# Prediction of plant organismal complexity based on transcription factor annotation: an AI approach

**DOI:** 10.64898/2026.08.18.745462

**Authors:** Deepti Varshney, Mustafa Hisham Tajjar, Jan de Vries, Frank Hutter, Stefan A. Rensing

## Abstract

How morphological complexity evolves is still enigmatic. While there is evidence in algae and plants as well as animals that diversification of the repertoire of transcription factors (TF) is causative for evolution of organismal complexity, there are many examples from lineages that follow their own way of complexity evolution, for example by expansion of particular families. For land plants, correlation of the size of the TF complement with number of cell types (as a proxy for morphological complexity) has been shown, and several families were identified as candidates to drive complexity evolution. Here, we expand a previously available dataset of cell type numbers from 12 to 82 proteomes and introduce a four class body plan scheme. We find that the total TF complement correlates with the number of cell types of Archaeplastida (primary plastid bearing plants and algae). We used TabPFN (Tabular Prior-data Fitted Network) for binary (uni- vs. multicellularity) as well as for four class *Bauplan* classification. TabPFN is able to predict the morphological complexity with high accuracy. This approach allows to determine organismal complexity based on the gene space of an organism. Based on our results, we can confirm that plant morphological evolution is driven by gain and expansion of TF families.

## Introduction

Complexity of organisms is often measured by the number of discrete (morphologically distinguishable) cell types that an organism consists of. However, cell type estimates are not easy to come by, in particular because cell types might not differ morphologically but by their molecular (transcriptomic/epigenetic) state. Transcription associated-proteins (TAPs) are a means to overcome cell type estimates. They comprise i) transcription factors (TF) that promote or repress transcription by sequence-specific binding to the DNA, ii) transcriptional regulators (TR) that act on gene regulation via protein-protein interaction or epigenetic modification and iii) putative TAPs (PT) the function of which still needs to be proven. Based on cell type estimates for 12 species, correlation of cell type number with total TAP and TF complement of the encoded genome was previously found for green plants (Lang et al., 2010).

There is a general tendency in plants to increase the TAP complement (both by gain of families and genes) concomitant with increasing morphological complexity. However, it should be noted that there are also deviating mechanisms. For example, the expansion of the C2H2 family in species as diverse as octopus and red algae has been argued to be instrumental in increasing complexity (Petroll et al., 2021). In red algae, expansion of the C2H2 family occurred after an evolutionary bottleneck, adaptation to extreme environments, that let to genome reduction. At the base of the red algal clade in which multicellularity evolved, and concomitant with its further evolution, C2H2 expanded. In the streptophyte alga *Chara braunii*, on the other hand, the trihelix family is significantly expanded and has been argued to have been instrumental for the secondary rise of complexity in this lineage (Nishiyama et al., 2018). For green algae in particular, “If there is one universal principle that emerges from comparing the evolution of multicellularity across green algal taxa, it is that there are no universal principles (Umen & Herron, 2021).” Hence, variations in gene presence/absence patterns are to be expected.

Terrestrialization of land by plants was enabled by evolutionary novelties such as abiotic and biotic stress tolerance, a sturdy cell wall with its peculiar phragmoplast-based mode of cell division, or diversified signaling pathways. On land, further novelties such as vasculature or stomata evolved. Underlying these morphological changes towards higher organismal complexity are additions to the TAP repertoire: there are in general more TAPs in streptophyte algae than in red and green algae, and more again in Embryophyta than in streptophyte algae (Petroll et al., 2025). While most plant-specific families were gained already in streptophyte algae, before the water-to-land transition, many families have gained members since (Bowman et al., 2017; Feng et al., 2024; Jiao et al., 2020; Lang et al., 2010; Petroll et al., 2025; Wilhelmsson et al., 2017). Acquisition of families includes horizontal gene transfer; for example, the TF GRAS family has been detected to be gained from bacterial progenitors in the last common ancestor (LCA) of Anydrophyta (Zygnematophyceae and Embryophyta), and argued to have been instrumental in acquiring abiotic stress tolerance (Jiao et al., 2020; Rensing, 2020).

While the coupling of TAP gain with increasing morphological complexity is true as a general trend, there are exceptions to the rule. For example, the Zygnematophyceae, sister lineage to land plants, have gone through a secondary loss of complexity (Feng et al., 2024; Goldbecker et al., 2026; Kunz et al., 2025), evident for example in their peculiar mode of sexual reproduction, devoid of motile sperm. While some species conserved this status, others like *Penium* or *Spirogloea* expanded their TAP repertoire again (Cheng et al., 2019; Jiao et al., 2020; Petroll et al., 2025). The same is true for *Chara*, as mentioned above. Indeed, likey *Chara*, many streptophyte lineages have likely gained again and again pronounced morphological complexity (Darienko et al., 2026), including the famous discoidal thalli of *Coloechaete scutate* (Bierenbroodspot et al., 2025). Some lineages of bryophytes also went through gene loss, most noteworthy in liverworts and hornworts (Harris et al., 2022; Li et al., 2020; Puttick et al., 2018). Mosses have also reduced their gene space from what was present in the LCA of Embryophyta, but expanded it again afterwards both by means of genome duplication and horizontal gene transfer (Dong et al., 2025; Petroll et al., 2025).

Here, we largely expanded previous datasets by means of adding additional genomes, and by annotating them with metadata about uni- or multicellularity, and their types of body plan. We estimate the number of cell types for species for which no estimates were available. Moreover, we use TabPFN (Hollmann et al., 2025), a pretrained transformer model for zero-shot predictions on tabular data, to classify body plan types based on TAP complements.

## Results and Discussion

### Correlation of total number of TAPs and TFs with the number of cell types

The pattern of overall numbers of TAPs as well as the numbers of TFs show positive correlation with the number of cell types as a measure for morphological complexity (Lang et al., 2010). Since that estimate was based on only 12 species, we added cell type estimates for 70 proteomes, bringing the number to 82 (Table S1). We calculated the best fit and generated a graph that contains both the known and estimated data for all TAPs and TFs only (Table S2/S3, File S1 and S2).

We find that the polynomial fit is a good approximation for the known data (all TAPs R^2^=0.73, TFs = 0.90). On average, absolute deviation between predicted and known is 67% for TAPs and 65% for TFs; for example, *Arabidopsis thaliana*: 27 “known” cell types, 34 predicted based on TAPs (27% deviation), 42 predicted based on TFs (54% deviation); or *Chlamydomonas reinhardtii*: 2 “known”, 2.9 based on TAPs (44%), 3.4 based on TFs (70%). There are no species where ≤100% fewer cell types were predicted, and there are 18/19 species for which ≥ 100% more cell types were predicted than “known”, for example *Cyanidium caldarium*: 2 “known”, 5.3 based on TAPs (163%), 4.7 based on TFs (136%). Most of these species are red algae, as well as a few streptophyte algae and a cryptophyte alga.

For the species with known cell type numbers the prediction error based on TAPs was 6.24 cell types (residual standard error). For TFs, the model was more accurate, with a typical deviation of 3.75 cell types, reflecting the higher R^2^ value (0.90) and better correlation between TFs and number of cell types.

Based on the fitted equation, we predicted the cell types for the remaining species whose numbers were previously unknown (Table S2/3). The prediction largely follows the expected taxonomic trends; Embryophyta generally show higher predicted cell type numbers compared to other taxonomic groups (File S1 and S2), reflecting their high morphological complexity. In contrast, streptophyte algae show a broader and more scattered distribution that may reflect both secondary loss and gain of complexity.

### Binary classification of uni- and multicellularity based on the TAP/TF complement

In order to analyse the complexity of Archaeplastida via means of their TAP complement (as measured by numbers of genes per family), we defined two categories for binary classification: class 0 (unicellular) and class 1 (multicellular) and assigned it to 40 (class 0) and 404 (class 1) species (Table S4 TAPs/S5 TFs). TabPFN (Tabular Prior-data Fitted Network) was fitted to assign the species to any of the binary classifications. Results from stratified five-fold cross-validation show strong agreement between the true and predicted labels. TabPFN correctly classified 400 species in class 1 with average precision (AP) = 1.0 and 36 species in class 0 with AP = 0.92 (Fig. S1a). Only 8 species (4 for each class) were misclassified when using all TAP families (Table S4, Fig. S1b). When using only the TF complement, 401 in class 1 (AP = 1.0) and 34 in class 0 (AP = 0.95) (Fig. S1c) were correctly classified, with 9 misclassified species (6 from class 0 predicted as class 1 and 3 from class 1 predicted as class 0) (Fig. S1d). Among both the analyses, the same seven species were misclassified (Table S4 & S5, Fig. S2 - S3). All species consistently mis-classified as multicellular, although they are unicellular, are non-green algae (Rhodophyta and Glaucophyta), while all species consistently classified as unicellular, although they are multicellular, are Chlorophyta. Most of the species mis-classified here show the expected trend (more or less cell types) based on the correlation analyses using TAP/TF numbers as well. For example, *Cyanidium caldarium* (Rhodophyta) is known to be unicellular with two cell types (vegetative/sexual stage). It is classified as multicellular based on the TAP as well as the TF complement, and predicted to have 5 (TAP) or 4 (TF) cell types based on the correlation.

### Complexity (body plan) classification based on the TAP/TF complement

Subsequently, we further defined four organisational categories (body plans) expected to coincide with rising complexity: simple, filamentous, thallus and body. Simple (1 to 3 cell types) refers to unicellular or colonial organisms with at most one more cell type than vegetative cells and zygote, while filamentous (3 to 7 cell types) refers to filamentous or colonial organisation with at least two cell types more than vegetative and zygote. Thallus (4 to ca. 10 cell types) can be sheet or body (thallus), syncytial or parenchymous organization with several cell types, while body (>= 20 cell types) refers to 3D multicellular body organization with cell-cell connections and several organs. We added these categories to a range of organisms (Table S6 TAPs and S7 TFs), and used this as training data for prediction of the four classes by TabPFN.

In this *Bauplan* classification, we were able to classify for the most part with similar accuracy as in the binary classification. The body category showed the highest accuracy with more than 99% correct classifications. The simple (90% correctness for TAPs, 95% for TFs) and filamentous (83% TAPs/73% TFs) classes were also predicted with high accuracy (Table S6/S7). In contrast, prediction for the thallus class showed more variability (accuracy 60% for TAPs, 50% for TFs).

These trends were further supported by precision-recall (PR) curves. High average precision and recall values were observed for body (TAPs & TFs AP = 1.00), simple (TAPs AP = 0.94; TFs AP = 0.95) and filamentous (TAPs AP = 0.93; TFs AP = 90) classes, while the thallus class showed lower performance (TAPs AP = 0.73; TFs AP = 0.63) (Fig 1). Overall, most *Bauplan* class categories were classified reliably, with the thallus class displaying difficulty in separation from other classes.

**Fig. 1.**
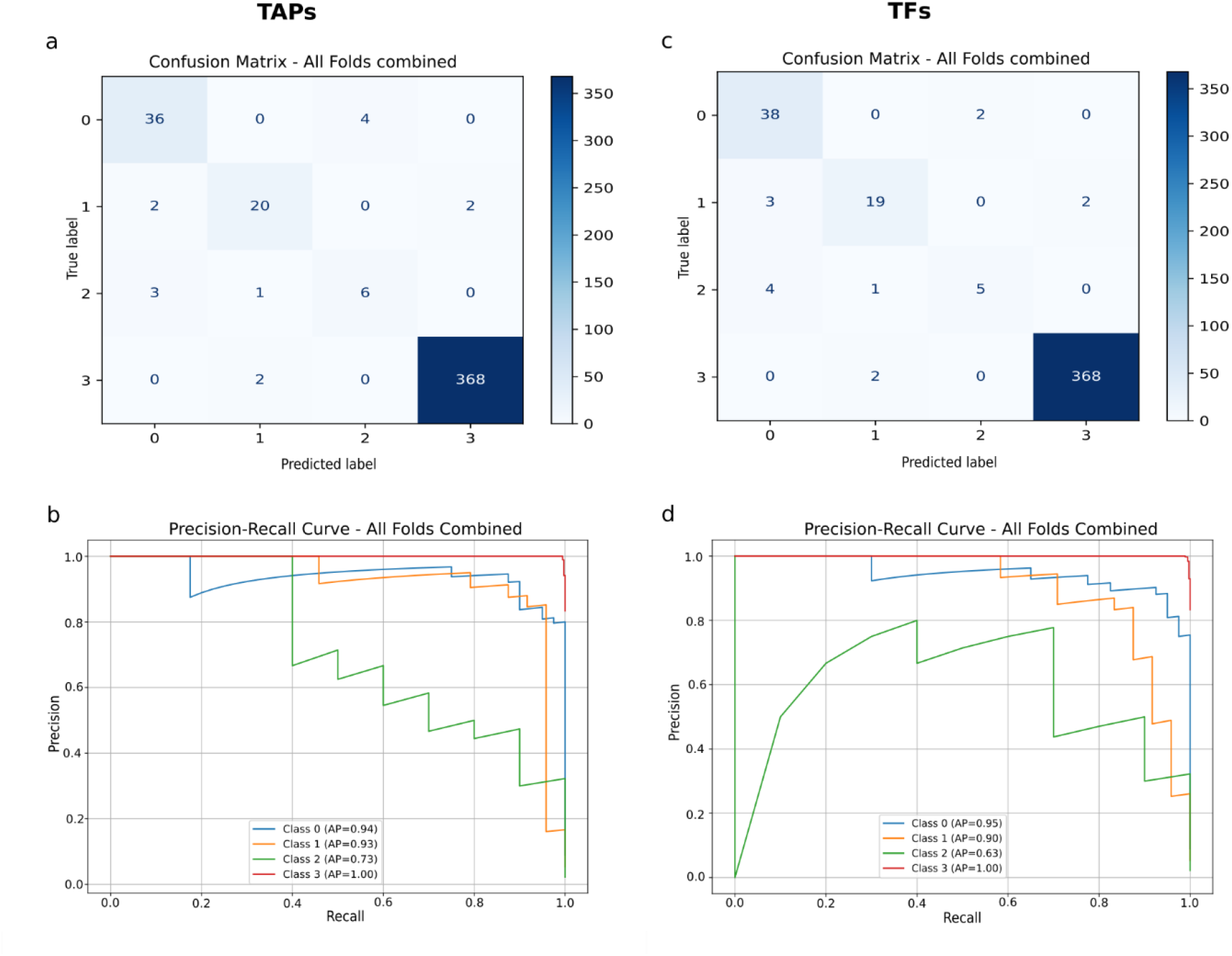
Evaluation of the TabPFN four-class *Bauplan* classification using the aggregated confusion matrix and class-specific precision-recall curves for TAPs (a, b) and TFs (c, d).

### Five-fold cross-validation consistency across folds and misclassification patterns

To evaluate classification performance, we examined the precision-recall curves across all five folds of stratified cross validation for both TAPs and TFs. Overall, PR curves show consistently high performances across folds for most categories (Fig. S6). For *Bauplan* classification, the body category (class 3) is classified almost perfectly in all folds with average precision 1.0 for both TAPs and TFs, indicating strong separability. The simple category (class 0) and filamentous (class 1) categories also show high and relatively stable performance across folds, with minor variation.

In contrast, the thallus category (class 2) shows lower and more variable performances across folds. While moderate performance is observed in folds 1-2, a significant drop in precision can be seen in folds 3-5 (Fig. S4-5; Table S10-13) indicating less stability compared to other classes. This pattern is consistent for both, TAPs and TFs. This difference can be explained by the strong class imbalance with the body class dominating (370 species), while the thallus category contains 10 species only. This small number of samples may be suboptimal for learning and classification.

To further understand the classification errors, we analyzed the misclassified species in the *Bauplan* classification. In total, 18 species were classified different from their known value for TAPs and TFs (Table S6/7). Among these, 10 species were misclassified in both datasets, while the remaining species are incorrectly classified in either the TAP or TF dataset. Misclassifications were observed across all categories; however they are not evenly distributed: the thallus category showed the highest error rate with 6 out of 10 species misclassified (60%). Species in this category were predicted as simple or filamentous categories, indicating a predictive shift towards lower morphological complexity. This pattern can be seen in chlorophyte species such as *Ulva mutabilis, Caulerpa lentillifera* and *Coleochaete sp*. as well as Rhodophyta species such as *Pyropia yezoensis, Porphyra umbilicalis and Calliarthron tuberculosum*. In the other *Bauplan* categories, misclassifications were less frequent and more variable. For example, *Volvox carteri* and *Mougeotia scalaris* were occasionally assigned to simple or body categories, while “body” category species such as *Funaria hygrometrica* and *Nitella hyalina* were predicted as “filamentous”.

Furthermore, we also compared SHAP (SHapley Additive exPlanations)-derived feature importance for all the species (correctly classified and misclassified) with the misclassified species in both TAP and TF datasets (Fig. 2). In both datasets, several gene families (features) showed higher importance in misclassified samples as compared to their overall contribution (Table S10/11). Comparison of the top 10 features of the misclassified species revealed that many of these were common between overall and misclassified samples, but with noticeably higher SHAP values in the misclassified set. For example, TAP families such as Trihelix, LFY, ARF, HD-NDX, TEA and CudA were among the most influential features in general, as well as among the misclassified species (Table S10/11). The increased importance of these features in misclassified species (as evidenced by their higher relative SHAP value) suggests that although they are important for prediction, they are not specific to a single *Bauplan* category. Overall, the SHAP analysis shows that misclassification is mainly caused by features that are important, but not specific enough to distinguish between *Bauplan* categories. To assess whether feature contributions remained stable across cross-validation folds, we compared SHAP importance across the individual folds. Several top 10 ranked features, including Trihelix, LFY, TEA, ARF, DUF246 and RF-X were consistently identified as top 15 ranking across all folds, with minor difference in feature ranking in both TAP- and TF-based predictions (Table S12/13). Hence, the major contributing features remained largely stable across cross-validation folds, while a few of the top features showed variation, for example Rel and CudA rank 135 and 136 in fold 5 (TAP-based prediction, Table S12) resp. 89 and 88 (TF-based prediction, Table S13).

**Fig. 2.**
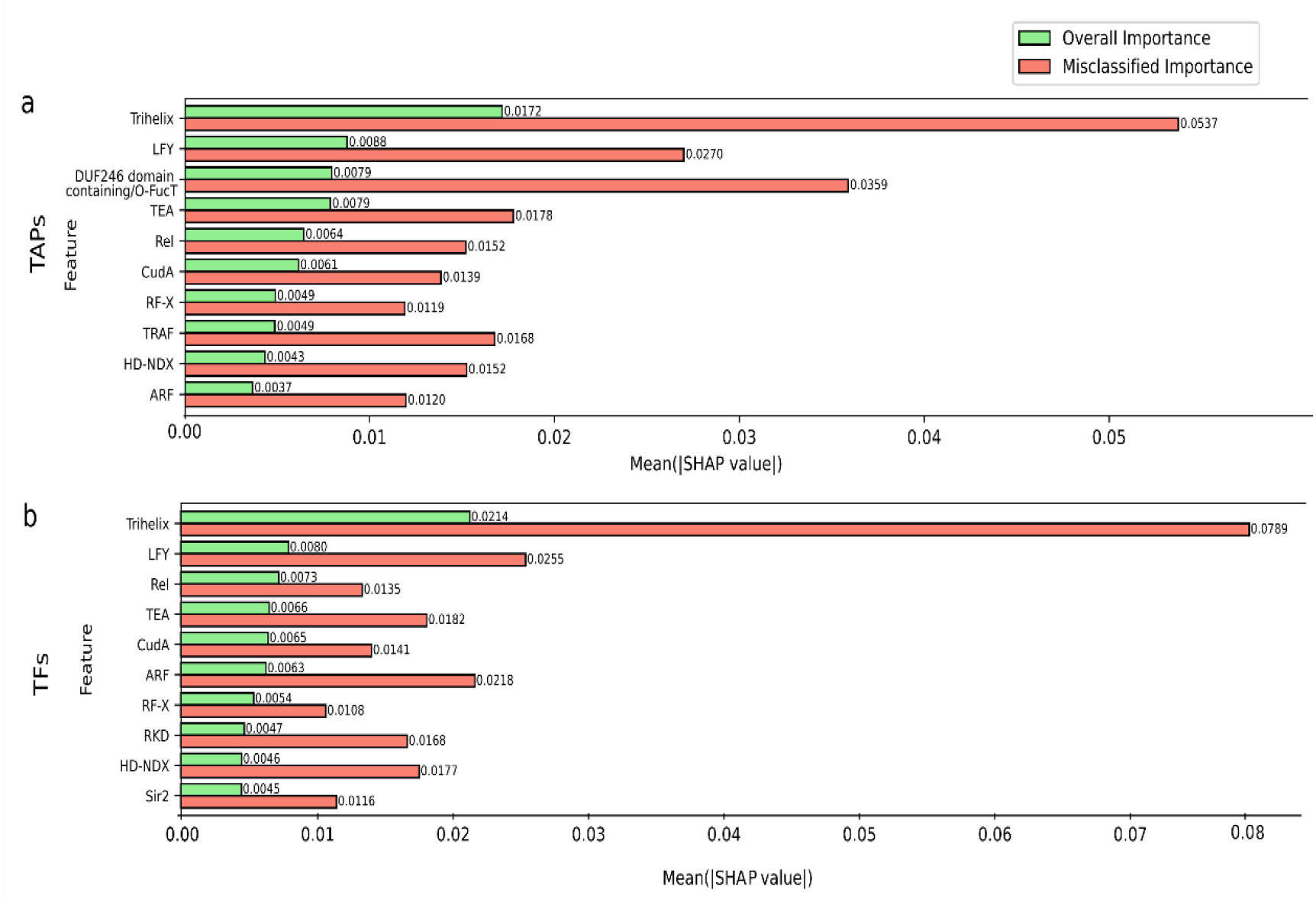
Top 10 SHAP-based features contributing to *Bauplan* classification based on TAP (a) and TF (b) complements. Mean absolute SHAP importance values averaged across five cross-validation folds are shown for overall (correctly classified and misclassified) vs. misclassified. Corresponding plots for all features are provided in Fig. S4/5.

### Caveats of classification approaches

All in all, only 18 of 444 species (4%) were classified different from the “known” (training) value in the multi-class (*Bauplan*) classification (Table S6/7). Of these 18, 14 were incorrectly classified using the TAP dataset, and 14 using the TF dataset, with an overlap of 10. Among these 18 species, 7 are represented by transcriptomic datasets, i.e. had no available genome-based gene annotation (Table S9). Transcriptome-based annotations are notoriously over-predicting gene space – hence we have the predictions for these species with a grain of salt.

If we consider the remaining 11 (2%), 7 are of non-green algae also found among the incorrectly classified species from the binary approach. From the remainder, 4 (all mis-classified in both approaches) are of the category multicellular green alga, with under-prediction of cell types/body plan complexity.

Together, the results indicate that TAP family number and composition provide a strong signal that correlates with morphological complexity as measured by number of cell types and body plan organization in Archaeplastida. We find that TFs alone provide a similar signal than all TAPs. This is in line with previous results (Lang et al., 2010) in the sense that the TF signal suffices for classification. We also find that while the methodology in general works well for Archaeplastida, there are restrictions to some lineages. Non-green species (Rhodophyta, Glaucophyta and Cryptophyta) are sometimes “over-predicted”, i.e. assigned a higher complexity than observed.

This might be due to lineage-specific expansion of certain gene families like C2H2 or HSF (Petroll et al., 2021).

Also, some multicellular Chlorophyta are “under-predicted”, i.e. assigned a lower complexity than observed. These species have gained their particular flavor of complexity independently from the lineage evolving into land plants – apparently, this cannot be predicted with ease by using the TF complement alone.

The training data used was size biased, with the “thallus” category containing the lowest number of samples. This might contribute to the lower accuracy of classification of the thallus category.

### Importance of individual TAP families for complexity evolution

We determined the top 10 families (using SHAP values) that contribute to the binary and four-way classification (Table S8). Out of the top 10 features that contribute to the uni- vs. multicellularity classification, there are four that previously have been hypothesized to contribute to multicellularity based on a principal component analysis (Lang & Rensing, 2015). Six of those found using the binary approach are also picked up among the top 10 for one or the other of the four classes. If we consider the top 10 contributing features for each of the four classes, there are two for class “simple” that were previously hypothesized to be involved in multicellularity, four for class “filamentous”, and three each for “thallus” and “body” (Fig. 3a). Eight families in total overlap between old and new approach (ARF, Aux/IAA, bZIP, C2C2_CO-like, C2H2, DUF246 domain containing/O-FucT, HD-NDX, TRAF) (Petroll et al., 2025)..

**Fig. 3.**
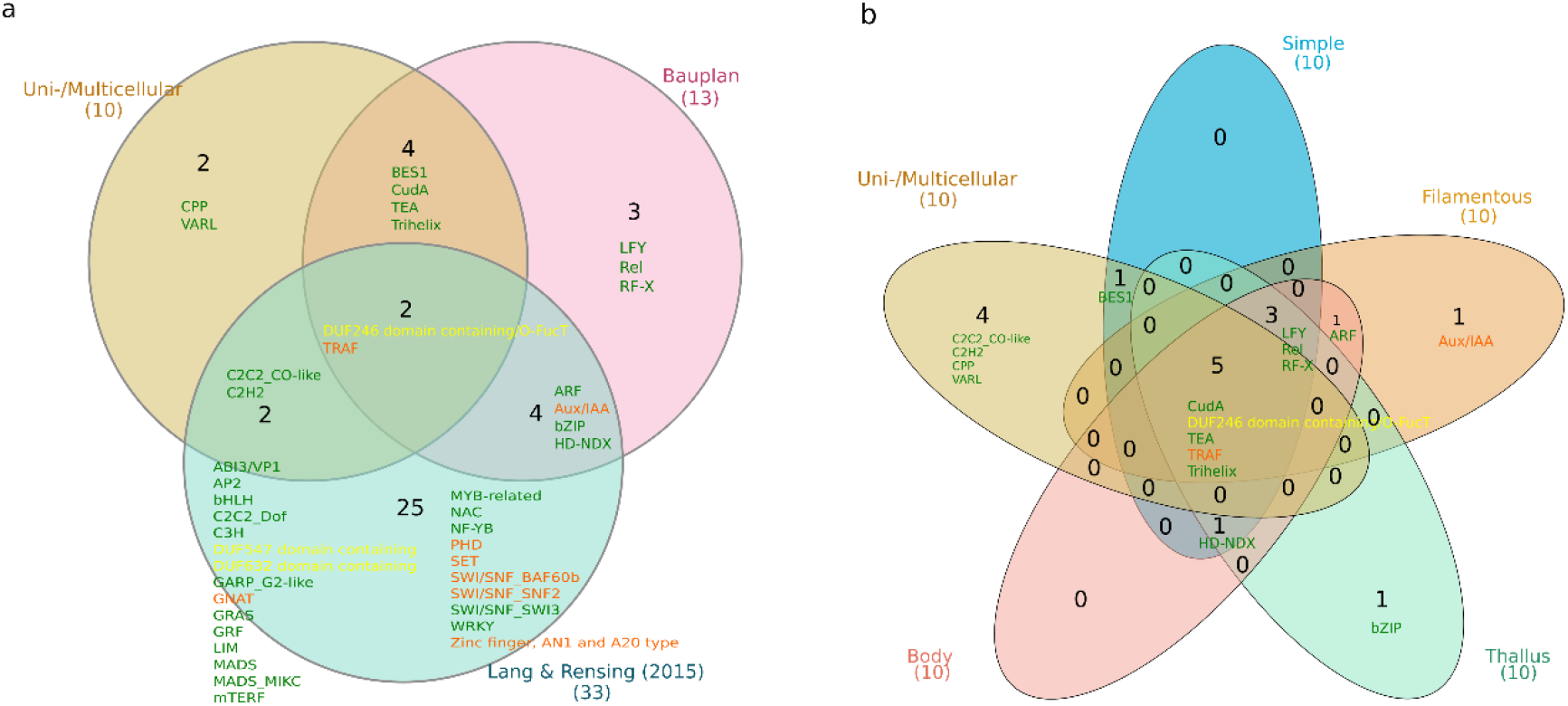
(a) Intersection of TAP families contributing to complexity based on a previous study and the present classification approaches, (b) Intersection of TAP families contributing to classification of organismal complexity in the present study.

Interestingly, there are also a couple of families that were not previously discussed and that contribute to cellularity classification (CPP, VARL), to both classifications (BES1, CudA, TEA, Trihelix) or the body organization classification (LFY, Rel, RF-X).

Strikingly, out of the 17 families that were found to contribute to classification, 14 are TF. There are two TR families, Aux/IAA and TRAF and one PT, DUF246 domain containing/O-FucT, that contribute. This latter family was identified as putative TAP DUF246 domain containing in (Richardt et al., 2007). Later, the defining domain was identified as O-FucT (GDP-fucose protein O-fucosyltransferase, PF10250) (Hansen et al., 2012). Decoration of proteins with O-Fucose by O-FucT has been shown to affect signal transduction in animals (Loriol et al., 2006) as well as influencing the plant male germ line via action on arabinogalactan synthesis (Stonebloom et al., 2016); O-fucosylation of DELLA activates DELLA-dependent plant development (Zentella et al., 2017). Our classification results reinforce the notion of the O-FucT family as a transcriptional regulator.

Many of the 17 features contribute to classification of several *Bauplan* classes; exceptions are ARF (“filamentous”, “body”) Aux/IAA (“filamentous”), Bes1 (“simple”) and bZIP (“thallus”) (Fig. 3b). HD-NDX contributes to all classes but filamentous; the families that contribute to classification of all features are CudA, DUF246 domain containing/O-FucT, LFY, Rel, RF-X, TEA, TRAF and Trihelix.

Among the 17 families detected by the present approach to be predictive for complexity (Table S8), 11 is known to act as homo-as well as heterodimers, while two are considered homodimeric only (DUF246 and LFY), one monomeric (CudA), while for two no data was found. TF homodimers are potentially more stable with regard to gene duplications, as their (homodimeric) interaction is not affected by dosage sensitivity of one of the two partner proteins. If TFs are able to act as heterodimers on top of the homodimeric interaction, as is the case for the above-mentioned 11 families, their combinatorial complexity potential increases. They are thus candidates for alteration of gene regulatory networks by duplication/retention and subsequent sub- or neofunctionalization of genes encoding the potential dimerization partners (Lang & Rensing, 2015). This has also been hypothesized e.g. for the plant MADS TF family (Veron et al., 2007). It is striking that 64% of the TAP families predictive for complexity may act as homo-as well as heterodimers, reinforcing the notion that they represent a playground for post-duplication gene regulatory network refinement.

Of the 19 families detected by the TabPFN approach, the majority (11, 58%) were gained at some point during Archaeplastida evolution (Table S8). Gain of TAP families and subsequent expansion and rewiring of gene regulatory networks hence contribute to evolution of morphological complexity.

## Conclusions

Classification of Archaeplastida organismal complexity based on tabular data on TAP/TF complements with the TabPFN foundation model ís generally possible with high accuracy. Genome-wide annotation of the TAP complement can easily be achieved using tools such as TAPscan. Given the constantly increasing number of genomes becoming available this methodology is a valid option for predicting cell type number and *Bauplan* of organisms. This is particularly interesting for lineages that are not well studied and allows quick and accurate assessment of complexity.

The majority of TAP families detected to be predictive for organismal complexity were gained during Archaeplastida evolution (58%), and are known to be able to act as homo-as well as heterodimers (64%). Together, this evidence suggests that gain of TAP families - in particular TF families (82%) - as well as the rewiring of gene regulatory networks after gene duplication events are causative for plant morphological evolution.

## Methods

### Dataset and cell type regression

The species dataset used in this study focuses on Chlorophyta and streptophyte algae, including non-seed plants and several Embryophyta (Table S1). Sequence data for all the species were obtained from Genome Zoo database (Petroll et al., 2025). To identify the number of TAPs/TFs gene families in all the species TAPscan (Petroll et al., 2025) v4 was utilized. To examine the relationship between the number of TAPs/TFs and number of cell types, we first extended the dataset of the number of cell types per species used in (Lang et al., 2010) from 12 to 73 (Table S1). Using the expanded dataset, we evaluated multiple regression approaches, including simple linear regression, logarithmic models, and polynomial regression. All statistical analyses were conducted in R and model fitting was performed using the *lm()* function. The calculation was done using 73 species with based on total numbers of TAPs/TFs per species and “known” number of cell types (Table S1-S3). Regression model fit was assessed using the coefficient of determination (R^2^); the best model in both cases was polynomial. The regression function was applied to predict the cell type numbers for the species for which cell types were unknown (Table S2/3). All visualizations were generated using the ggplot2 package and plotly/htmlwidgets packages to generate an interactive configurable HTML figure in R version 4.3.1 (Team, 2023) (Supplementary File 1&2).

### AI-based classification using TabPFN

To analyse the morphological complexity of Archaeplastida using their TAP/TF complement (as measured by numbers of genes per family), we classified all the species into two categories: unicellular (class 0) and multicellular (class1) (Table S4 TAPs/S5 TFs). Multicellular species were further subdivided into four classes reflecting increasing morphological complexity: simple, filamentous, thallus and body (class 0-3) (Table S6 TAPs/S7 TFs). For classification, we employed TabPFN v2.5 (Hollmann et al., 2025), a pretrained transformer-based architecture designed for tabular data. To ensure a robust and unbiased evaluation of the model, a stratified 5-fold cross-validation approach was implemented so that the relative frequency of each class is preserved in every split (Fig S6). The TabPFN classifier was then fitted on each fold, with GPU acceleration being used to speed up the process. Performance was measured by examining confusion matrices and precision-recall curves for each class, along with their average precision scores. All the calculations were conducted using the TabPFN implementation in Python (Table S4/5).

### LLM-based quaternary structure literature mining

To determine whether members of the 17 TAP families found to contribute to the “Bauplan” classification act as monomers, homodimers, heterodimers or combinations thereof, we conducted an LLM-augmented literature analysis using Claude Sonnet 4.5 (Anthropic, 2024) via the claude.ai web interface. For each family, the model was prompted to search the web and summarize published evidence on quaternary structure (Table S14). The prompt was as follows: “Conduct a web search to retrieve literature about the quaternary protein structure of the transcription factor “queried family”. Scan the results to determine whether queried family proteins are described to act as monomers, homodimers, heterodimers or homo- and heterodimers”.

## Data availability

All the sequence fasta files for all the gene spaces used in this study are available on MAdLandDB/Genome Zoo (https://github.com/Rensing-Lab/Genome-Zoo).

## Acknowledgements

This project was carried out in the framework of MAdLand (https://madland.science, DFG priority programme 2237), SAR is grateful for funding by the DFG (RE 1697/22-1, project ID 527529748). Mustafa Hisham Tajjar is supported by the Konrad Zuse School of Excellence in Learning and Intelligent Systems (ELIZA; https://eliza.school/) through the DAAD programme “Konrad Zuse Schools of Excellence in Artificial Intelligence,” sponsored by the Federal Ministry of Education and Research. FH would like to acknowledge funding by the DFG CRC SmallData (SFB 1597), grant number 499552394.

