## Supplementary material for "Prediction of plant organismal complexity based on transcription factor annotation: an AI approach": Fig. S1-S6

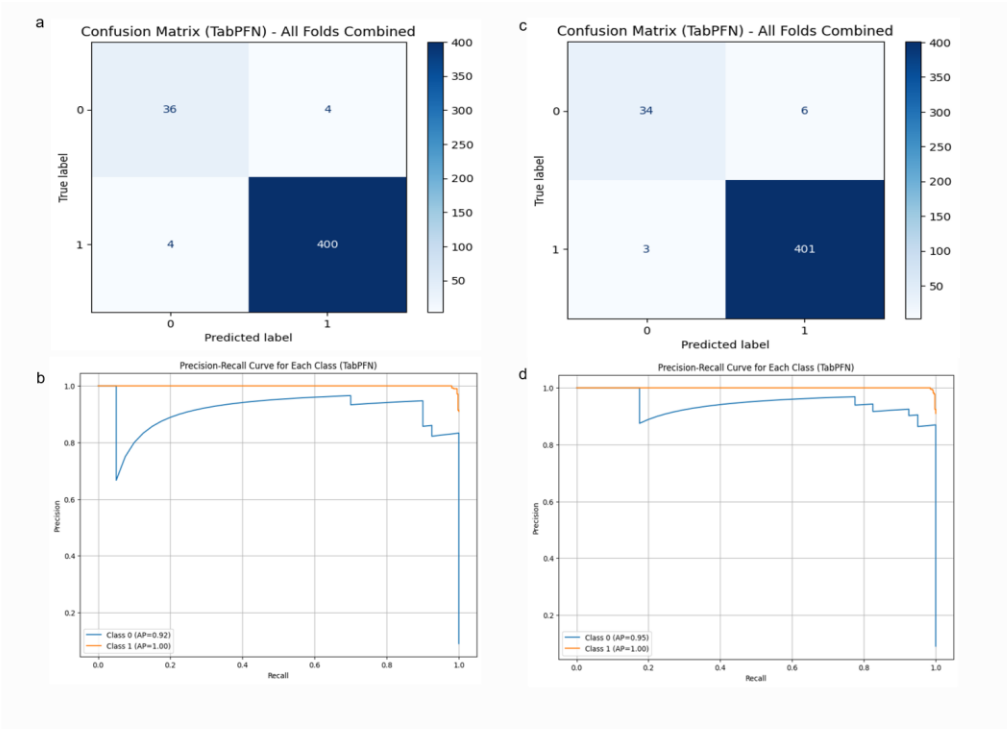


Fig S1. Evaluation of the TabPFN classifier using the aggregated confusion matrix and class-specific precision-recall curves for TAPs (a, b) and TFs (c, d) for binary classification.

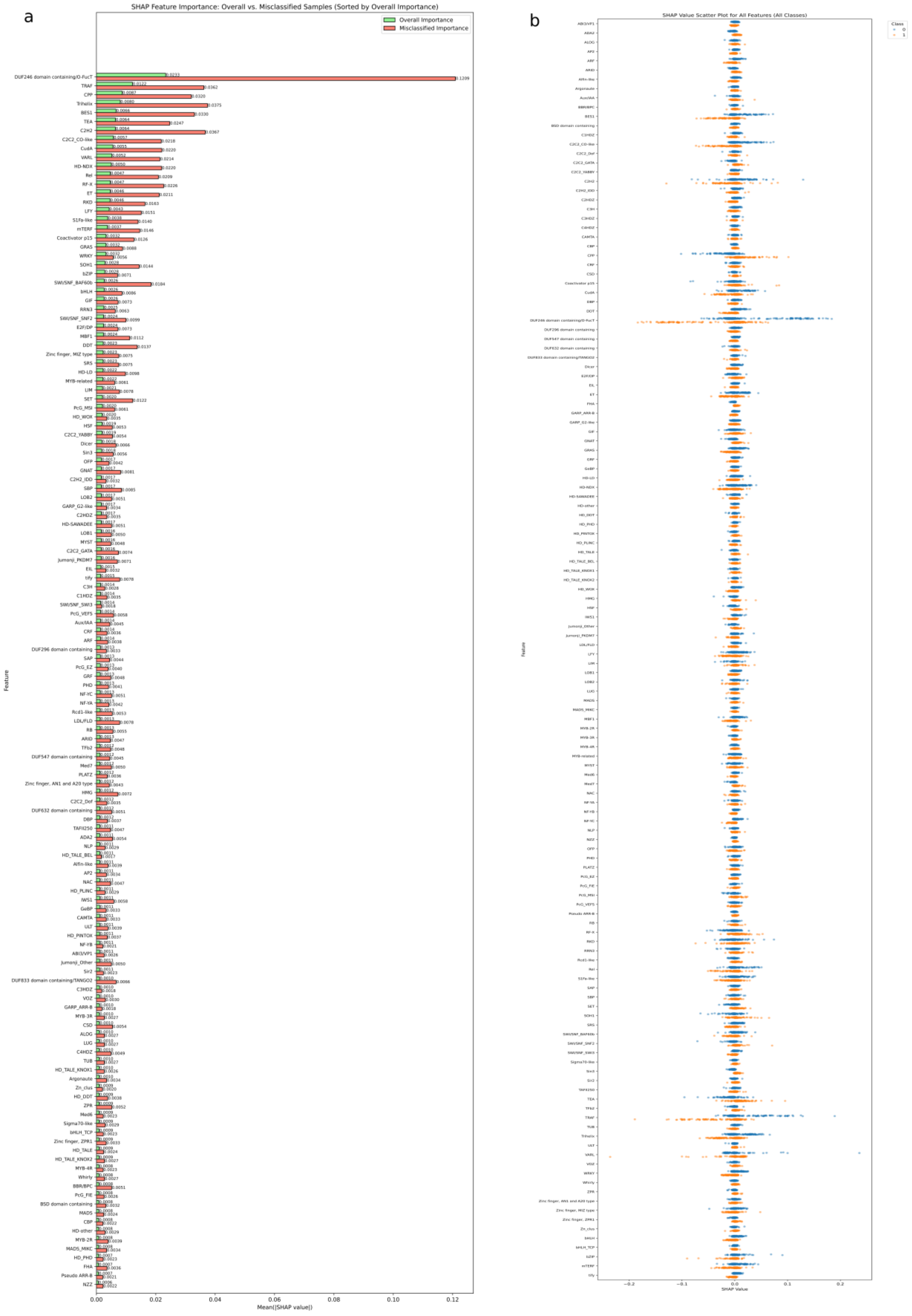


Fig S2. SHAP-based feature importance in TAPs binary classification (a) Average absolute SHAP feature importance across folds for overall (correctly +misclassified) and misclassified; (b) SHAP scatter plot showing the distribution of feature contributions across all classes.

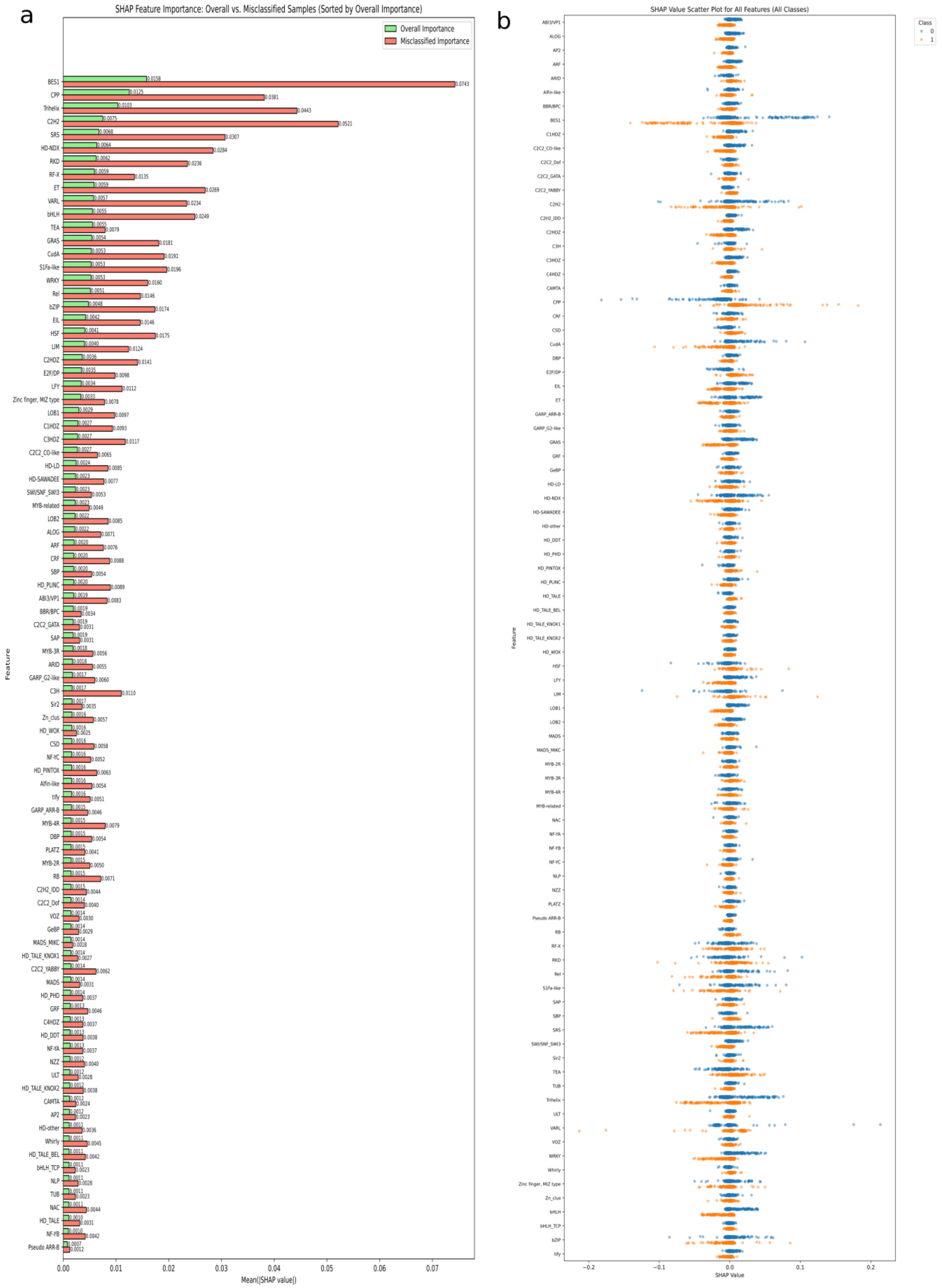


Fig S3. SHAP-based feature importance in TFs binary classification (a) Average absolute SHAP feature importance across folds for overall (correctly +misclassified) and misclassified; (b) SHAP scatter plot showing the distribution of feature contributions across all classes.

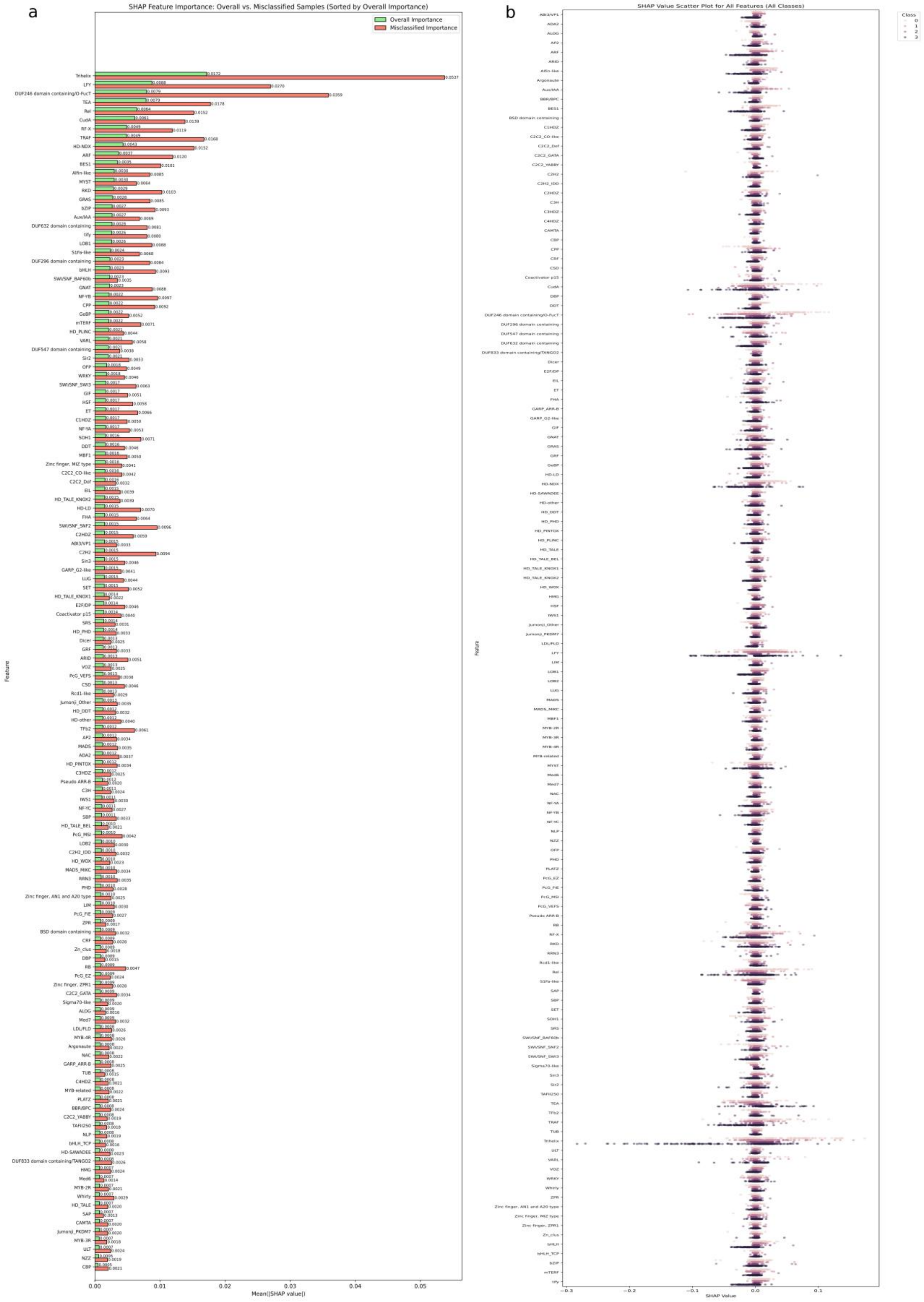


Fig S4. SHAP-based feature importance in TAPs *Bauplan* classification (a) Average absolute SHAP feature importance across folds for overall (correctly + misclassified) and misclassified; (b) SHAP scatter plot showing the distribution of feature contributions across all classes.

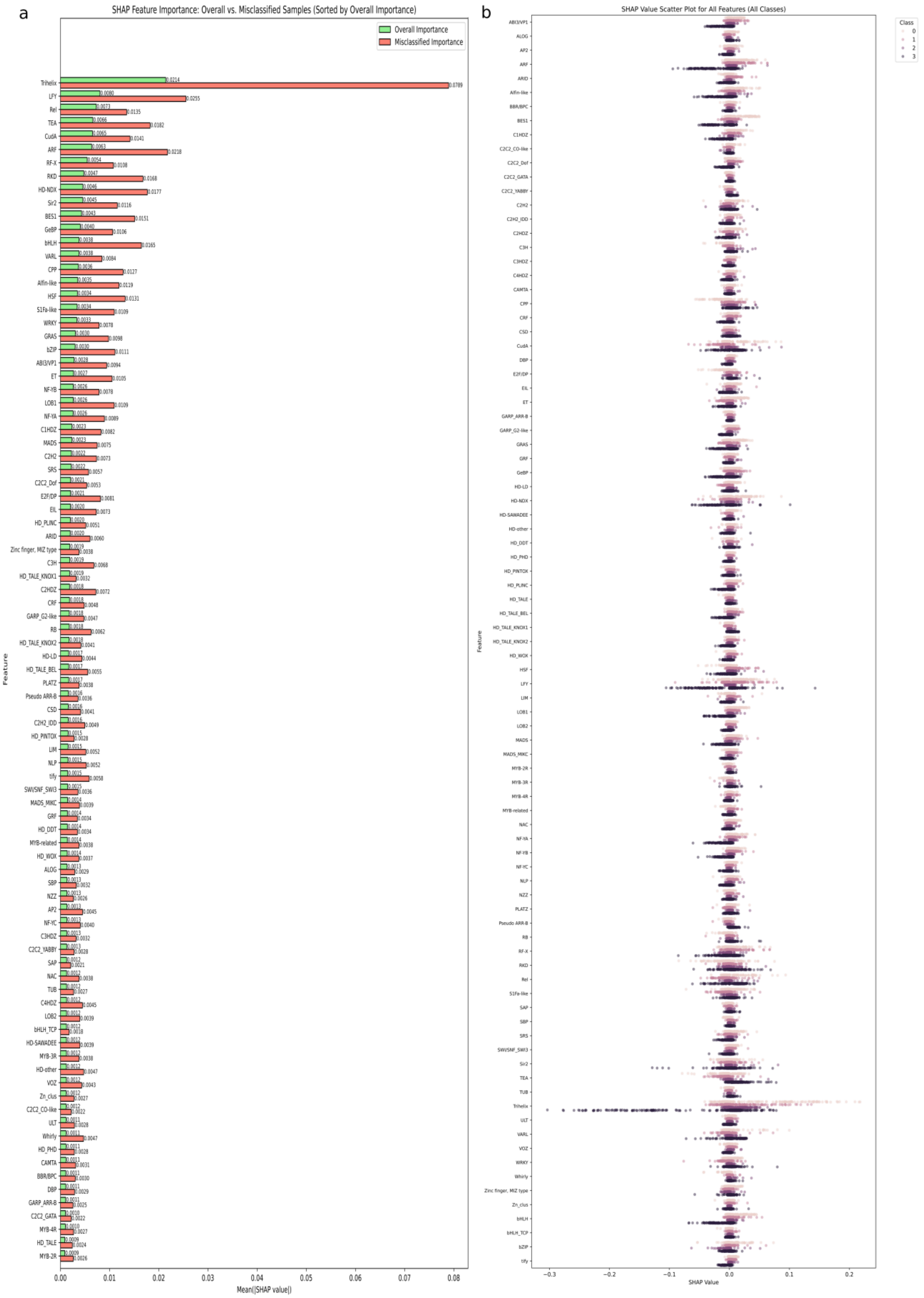


Fig S5. SHAP-based feature importance in TFs *Bauplan* classification (a) Average absolute SHAP feature importance across folds for overall (correctly + misclassified) and misclassified;(b) SHAP scatter plot showing the distribution of feature contributions across all classes.

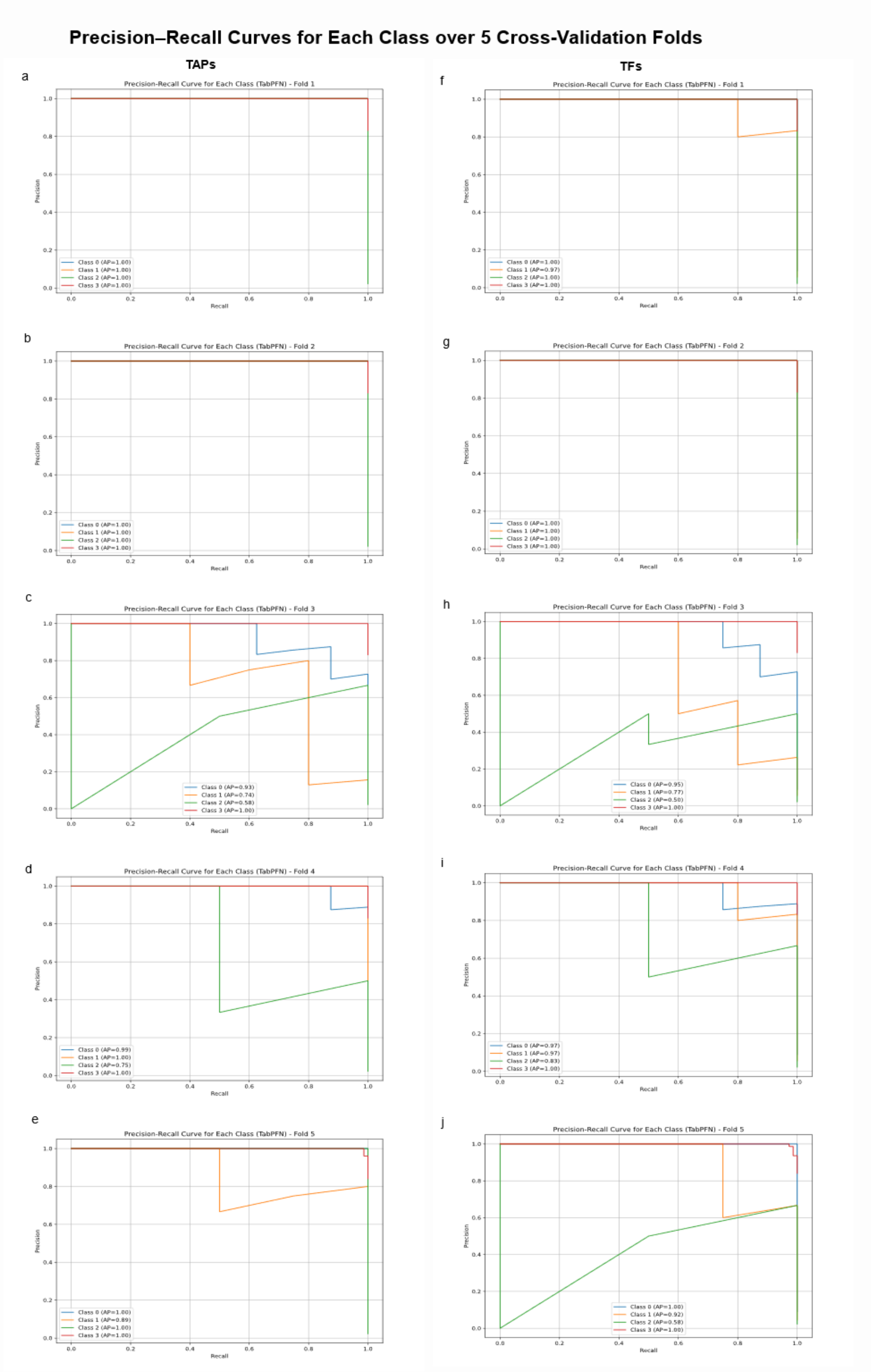


Fig S6. Evaluation of the TabPFN classifier using class-specific precision–recall curves across 5-fold cross-validation For TAPs (a-e, left panel) and TFs (f-j, right panel): *Bauplan* classification.
